# Functional conservation of Anopheline linalool receptors through 100 million years of evolution

**DOI:** 10.1101/2022.09.19.508609

**Authors:** Robert M. Huff, R. Jason Pitts

## Abstract

Insects rely on olfactory receptors to detect and respond to diverse environmental chemical cues. Detection of semiochemicals by these receptors modulates insect behavior and has a direct impact on species fitness. Volatile organic compounds (VOCs) are released by animals and plants and can provide contextual cues that a blood meal host or nectar source is present. One such VOC is linalool, an enantiomeric monoterpene, that is emitted from plants and bacteria species. This compound exists in nature as one of two possible stereoisomers, (*R*)-(–)-linalool or (*S*)-(+)-linalool. In this study, we use a heterologous expression system to demonstrate differential responsiveness of a pair of Anopheline odorant receptors (Ors) to enantiomers of linalool. The mosquitoes *Anopheles gambiae* and *Anopheles stephensi* encode single copies of Or29 and Or53, which are expressed in the labella of *An. gambiae*. (*S*)-(+)-linalool activates Or29 orthologs with a higher potency than (*R*)-(–)-linalool, while the converse is observed for Or53 orthologs. The conservation of these receptors across a broad range of Anopheline species suggests they may function in the discrimination of linalool stereoisomers, thereby influencing the chemical ecology of mosquitoes. One potential application of this knowledge would be in the design of novel attractants or repellents to be used in integrated pest management practices.

## Introduction

Mosquitoes utilize peripheral chemosensory receptors to detect volatile organic compounds (VOCs) in their environments that have a direct influence on species fitness. Chemosensory receptor proteins are encoded by three large gene families in insects: odorant receptors (Ors), gustatory receptors (Grs), and variant ionotropic glutamate receptors (Irs) (Benton *et al*. 2009; Freeman *et al*. 2014; Robertson 2019; Suh *et al*. 2014). These receptors display a significant amount of divergence both within and across families (Benton *et al*. 2009; Freeman *et al*. 2014; Robertson 2019; Suh *et al*. 2014). Oviposition site selection and animal host seeking are exclusively female behaviors and are influenced by the detection of VOCs (Takkenberg *et al*. 1991; Clements 1992; Zwiebel and Takken 2004; Carey *et al*. 2010; McBride 2016). Odor detection also influences nectar feeding and resting site selection (Otienoburu *et al*. 2012; Chaiphongpachara *et al*. 2018). The molecular foundations of chemosensory biology remain poorly understood in most mosquito species. A more thorough characterization of the molecular interactions between chemosensory receptors and their cognate semiochemicals is essential for understanding mosquito behavior. Previous studies have contributed to our understanding of how odor cues can attract mosquitoes to suitable animal hosts and oviposition sites, as well as to nectar sources (Foster and Hancock 1994; Takken and Knols 1999; Bernier *et al*. 2000; Allan *et al*. 2005; Riffell 2011; McBride *et al*. 2014; Chen and Kearney 2015). Expanding our knowledge of how chemosensory receptors respond to both plant and animal VOCs is essential for a more complete understanding of insect chemical ecology.

VOCs are emitted from a number of flowering plants. One of the most ubiquitous constituents is a terpene alcohol, linalool, which exists as one of two stereoisomers, (R)-(–)-linalool or (S)-(+)-linalool (Pereira *et al*. 2018; Aprotosoaie *et al*. 2014). Linalool is also emanated from the surfaces of human skin in association with the composition of the skin microbiome and may play a role in blood-host selection by anthropophilic mosquito species (Logan *et al*. 2008; Roodt *et al*. 2018). Linalool is perceived by many animal taxa and is known to affect the behavior of insects generally (Pichersky *et al*. 1995; Pichersky and Gershenzon 2002). It can act as a pollinator attractant, oviposition stimulant, and spatial repellent (Pichersky *et al*. 1995; Jonsson and Anderson 1999; Kline *et al*. 2003; Malo *et al*. 2004; Rostelien *et al*. 2005; Ulland *et al*. 2006; Reisenman *et al*. 2010a; Yu *et al*. 2015). There is also evidence that insect chemosensory receptors are enantiomerically selective for linalool and might play important roles in the chemical ecologies of diverse species (Ulland *et al*. 2006; Reisenman *et al*. 2010a). Insect behaviors in response to the detection of sex pheromones often occur on a enantioselective basis with few defined ligand/receptor relationships. Structural isomers of the sex pheromone japonilure produces attractive or repulsive behavior in a species specific manner for coleopterans (Tumlinson *et al*. 1977; Leal 1996). In the silkmoth *Bombyx mori*, a male restricted enantioselective sex pheromone receptor, BmOR1 was identified to detect the pheromone bombykol (Sakurai *et al*. 2004). Enantioselectivity, or at least strong bias in stereoisomer responsiveness, of mosquito chemosensory receptors has been demonstrated previously (Bohbot and Dickens 2009; Dekel *et al*. 2016). For example, the strict nectar feeding mosquito *Toxorhynchites amboinensis* displays selectivity of Or8 to the (R)-enantiomer of octenol (Dekel *et al*. 2016).

We have previously identified linalool as a potent activator of *Anopheles gambiae* odorant receptor 29 (AgamOr29; Huff and Pitts 2019). AgamOr29 is one of only a handful of labellar expressed odorant receptors and has been shown to be highly expressed in both male and female labella in addition to low level antennal expression (Pitts *et al*. 2011; Rinker *et al*. 2013; Saveer *et al*. 2018). Determining the responsiveness of labellar odorant receptors will help in our understanding of how mosquitoes are influences by important environmental cues. In this study, we have used a cell-based functional assay to further characterize the responsiveness of AgamOr29 towards linalool stereoisomers and structurally similar terpenoid compounds. Further, we have shown that a paralogous receptor, AgamOr53, is tuned to both enantiomers of linalool with slight differences in potencies. We chose to characterize AgamOr53 due to the conservation of amino acid residues in relation to AgamOr29. Importantly, we also demonstrate that orthologous receptors in *Anopheles stephensi*, AsteOr29 and AsteOr53, are tuned to linalool with differential responsiveness to linalool isomers. *Anopheles stephensi* is an evolutionarily distinct lineage, separated by millions of years of evolution from *Anopheles gambiae*, despite this, the paralogs retain similar functionalities of linalool detection (Neafsey *et al*. 2015).The conservation of Or29 and Or53 across Anophelines suggests that this ecologically ubiquitous compound might serve as an attraction cue for nectar-seeking mosquitoes or as a blood meal host attractant. Moreover, these receptors could serve as targets for the development of new excito-repellents that are specific to malaria vectors.

## Materials and methods

### Phylogenetic analysis

*An. gambiae* and *An. stephensi* Or29 and Or53 homologs were identified in all available Anopheline genomes on NCBI via tBLASTn or BLASTp searches (29). New gene annotations or reannotations were performed by identifying homologous exon regions and canonical intron splice junctions, also taking into account conserved intron positions across species. We have presented conceptual translations for all gene annotations in Supplementary table S1. Microsyntenic relationships were identified using the same methodology. Geneious Prime^2019^ software (Biomatters Limited, USA) was utilized for multiple sequence alignment as well as phylogenetic tree construction using the Neighbor-Joining method with 1000 bootstrap pseudoreplicates. In order to better resolve relationships among Or29/53 homologs, a selected sample of unrelated *Anopheles gambiae* Ors were used to construct the alignment and tree, with AgamOrco serving as the outgroup.

### Gene cloning and sequencing

AgamOrco and AgamOr29 cRNAs were synthesized as described previously (Huff and Pitts 2019). AgamOr53, AsteOrco, AsteOr29, and AsteOr53 templates were synthesized by Twist Biosciences (San Francisco, CA, USA) and cloned into the pENTR^*TM*^ vector using the Gateway^*R*^ directional cloning system (Invitrogen Corp., Carlsbad, CA, USA) and subcloned into the *Xenopus laevis* expression destination vector pSP64t-RFA. Plasmids were purified using GeneJET Plasmid Miniprep Kit (ThermoFisher Scientific, Waltham, MA, USA) and sequenced in both directions to confirm complete coding regions.

### Chemical reagents

The chemicals (S2 Table) used for the deorphanization of receptors were obtained from Acros Organics (Morris, NJ, USA), Alfa Aesar (Ward Hill, MA, USA), ChemSpace (Monmouth Junction, NJ, USA), Sigma Aldrich (St. Louis, MO, USA), TCI America (Portland, OR, USA), and Thermo Fisher Scientific (Waltham, MA, USA) at the highest purity available (Supplementary Table S2). (S)-(+)-linalool was custom ordered and synthesized by ChemSpace (Monmouth Junction, NJ, USA). Or29/Or53 chemical class responsiveness was determined by using complex odorant blends comprising 119 unique chemical compounds. Odorants were made into 1M stocks in 100% DMSO. Compounds in blends were grouped by chemical class and were combined and diluted to 10^−4^ M in ND96 perfusion buffer (96mM NaCl, 2mM KCl, 5mM MgCl_2_, 0.8mM CaCl_2_, and 5mM HEPES).

### Two-electrode voltage clamp of *Xenopus laevis* oocytes

AgamOrco, AgamOr29, AgamOr53, AsteOrco, AsteOr29, and AsteOr53 cRNA were synthesized from linearized pSP64t expression vectors using the mMESSAGE mMACHINE^®^ SP6 kit (Life Technologies). Stage V-VII *Xenopus laevis* oocytes were ordered from Xenopus1 (Dexter, MI, USA) and incubated in ND96 incubation media (96 mM NaCl, 2mM KCl, 5mM HEPES, 1.8mM CaCl_2_, 1mM MgCl_2_, pH 7.6) supplemented with 5% dialyzed horse serum, 50 μg/mL tetracycline, 100μg/mL streptomycin, 100μg/mL penicillin, and 550 μg/mL sodium pyruvate. Oocytes were injected with 27.6 nL (27.6 ng of each cRNA) of RNA using the Nanoliter 2010 injector (World Precision Instruments, Inc., Sarasota, FL, USA). Odorant-induced currents of oocytes expressing functional odorant receptor complexes were recorded using the two-microelectrode voltage-clamp technique (TEVC). The OC-725C oocyte clamp (Warner Instruments, LLC, Hamden, CT, USA) maintained a -80mV holding potential. To measure the effect of the blends on complexes, we perfused 10^−4^ M concentration blends. Current was allowed to return to baseline between drug administrations. Data acquisition and analysis were carried out with the Digidata 1550 B digitizer and pCLAMP10 software (Molecular Devices, Sunnyvale, CA, USA). A tuning curve was generated using a panel of 9 odorants including (*R*)-(–)-linalool, (*S*)-(+)-linalool, and closely related structural odorants. All chemicals used were administered at 10^−4^ M. All data analyses were performed using GraphPad Prism 8 (GraphPad Software Inc., La Jolla, CA, USA). For the establishment of a concentration-response curves, oocytes were exposed to (*R*)-(–)-linalool or (*S*)-(+)-linalool (10^−7^ M to 10^−4^ M). To measure the effect of the compounds on the oocytes, odorants were perfused for 10s. Current was allowed to return to baseline between chemical compound administrations.

## Results

### Anopheline linalool receptor homologs

The genomes of *An. gambiae* and *An. stephensi* encode single copies of Or29 (AGAP009111, ASTE010099) and Or53 (AGAP009390, ASTE015875), with apparent 1:1 orthologs encoded in the genomes of several additional Anopheline species, totaling 19 in all (Figure 1A). We identified orthologs and paralogs via BLAST searches. Some genes were not annotated in the current genome assemblies, while others were incomplete or inaccurately annotated, based on amino acid alignment. In either of those instances, we annotated or reannotated genes to maximize overall amino acid homology and conserved intron positions. We have presented our conceptual translations in Supplementary Table S1. As shown in Figure 1A, Or29 and Or53 form distinct clades across a number of Anopheline species with high bootstrap support. Interestingly, Or29 orthologs were only identified within the subgenus, Cellia, while the subgenera Nyssorhynchus and Anopheles lack apparent orthologs (Figure 1A). This could be the result of gene gain in the former lineages or gene loss in the latter. Similarly, genes with strong homology, perhaps orthology, to Or53 were identified in the Cellia subgenus and paralogs, including Or30, were found across all subgenera (Figure 1A). Amino acid alignments for AgamOr29/AsteOr29 and AgamOr53/AsteOr53 were generated to demonstrate the level of identity and homology between lineages (Figure 1B). A clearly conserved gene structure of four exons and three introns was evident across nearly all Or29 and Or53 homologs (Figure 1C). Pairwise identities are 79% and 67% for Or29 and Or53, respectively, with conserved residues spanning their entire lengths (Figure 1D; Supplementary Table S1). The relationships across these two ORs conform to previously described Anopheline species relationships (Manguin 2013; Neafsey *et al*. 2015; Norris and Norris 2015). Insect odorant receptors are generally divergent, with evidence for rapid evolution and positive selection that is amongst the highest for major gene families in Anophelines (Neafsey *et al*. 2015). Despite this, a comparison of 45 potential one-to-one orthologs between *An. gambiae* and *An. stephensi* revealed an average amino acid identity of 73.5% with a standard deviation of 12.4% (Supplementary Table S4).

**Fig. 1.**
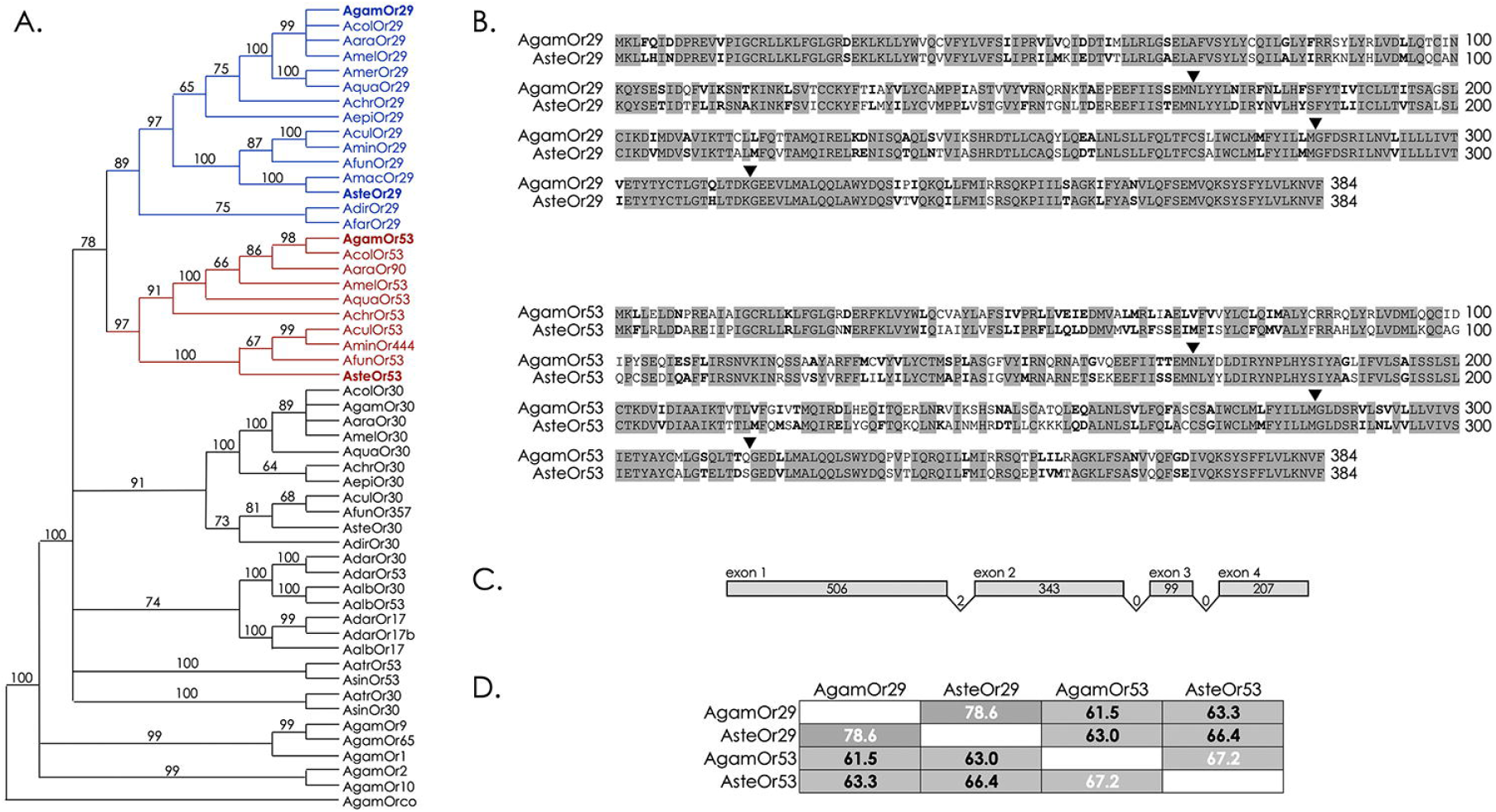
Conserved Anopheline Odorant Receptors. (A) Neighbor-joining tree showing relationships between anopheline Odorant receptor conceptual peptides. Node values signify bootstrap support from 1000 pseudoreplicates. Or29 clade shown in blue, Or53 clade shown in red. (B) Amino acid alignment of Or29 (Top) and Or53 (Bottom) orthologs from *Anopheles gambiae* and *An. stephensi*. Identical residues are highlighted in grey and similar residues are in bold type. Positions of introns are indicated with filled triangles. Amino acid numbering is indicated at the end of each line. (C) Conserved exon/Intron structure of Or29 and Or53 orthologs. Exons are shown as gray boxes with lengths given in base pairs. Introns phases between exons are indicated by numbers. (D) Table of amino acid identities across *An. gambiae* and *An. stephensi* odorant receptor homologs. Alphabetical species abbreviations: Aalb (*An. albimanus*), Aara (*An. arabiensis*), Aatr (*An. atroparvus*), Achr (*An. christyi*), Acol (*An. coluzzii*), Acul (*An. culicifacies*), Adar (*An. darlingi*), Adir (*An. dirus*), Aepi (*An. epiroticus*), Afar (*An. farauti*), Afun (*An. funestus*), Agam (*An. gambiae*), Amac (*An. maculatus*), Amel (*An. melas*), Amer (*An. merus*), Amin (*An. minimus*), Aqua (*An. quadriannulatus*), Asin (*An. sinensis*), Aste (*An. stephensi*).

### Conserved genetic structure and microsynteny of linalool receptor homologs

Observable patterns of chromosomal microsynteny further support the evolutionary relationships of the group of linalool receptors that we have proposed (Figure 2). Well conserved order and clustering of neighboring genes near linalool receptors supports preservation of chromosomal segments in Anophelinae and this has been observed in other species of Diptera (Pitts *et al*. 2021; Ruel *et al*. 2021). In certain cases, such as the Or29 cluster in *An. stephensi*, complete conservation of gene order was not apparent, possibly due to the limitation of the sizes and resolutions of the Anopheline genome assemblies (Figure 2). In addition, the transcriptional directions of the nearby orthologous genes is also overwhelmingly static. Or53 genetic regions within Anophelines encompass a cluster of odorant receptor genes (Figure 3) that are expressed as polycistronic messages (Karner *et al*. 2015). Interestingly, these genes seem to be expressed in the labellum of *An. coluzzii* (Saveer *et al*. 2018), comprising the majority of the Or expression in this appendage (Figure 4). An additional Or cluster (Or3,4,5) is also expressed in the labellum (Fox *et al*. 2002). The functionality of these receptors has been explored, in part, via heterologous expression, with activators representing various chemical classes (Carey *et al*. 2010; Wang *et al*. 2010). However, linalool was not among the identified ligands.

**Fig. 2.**
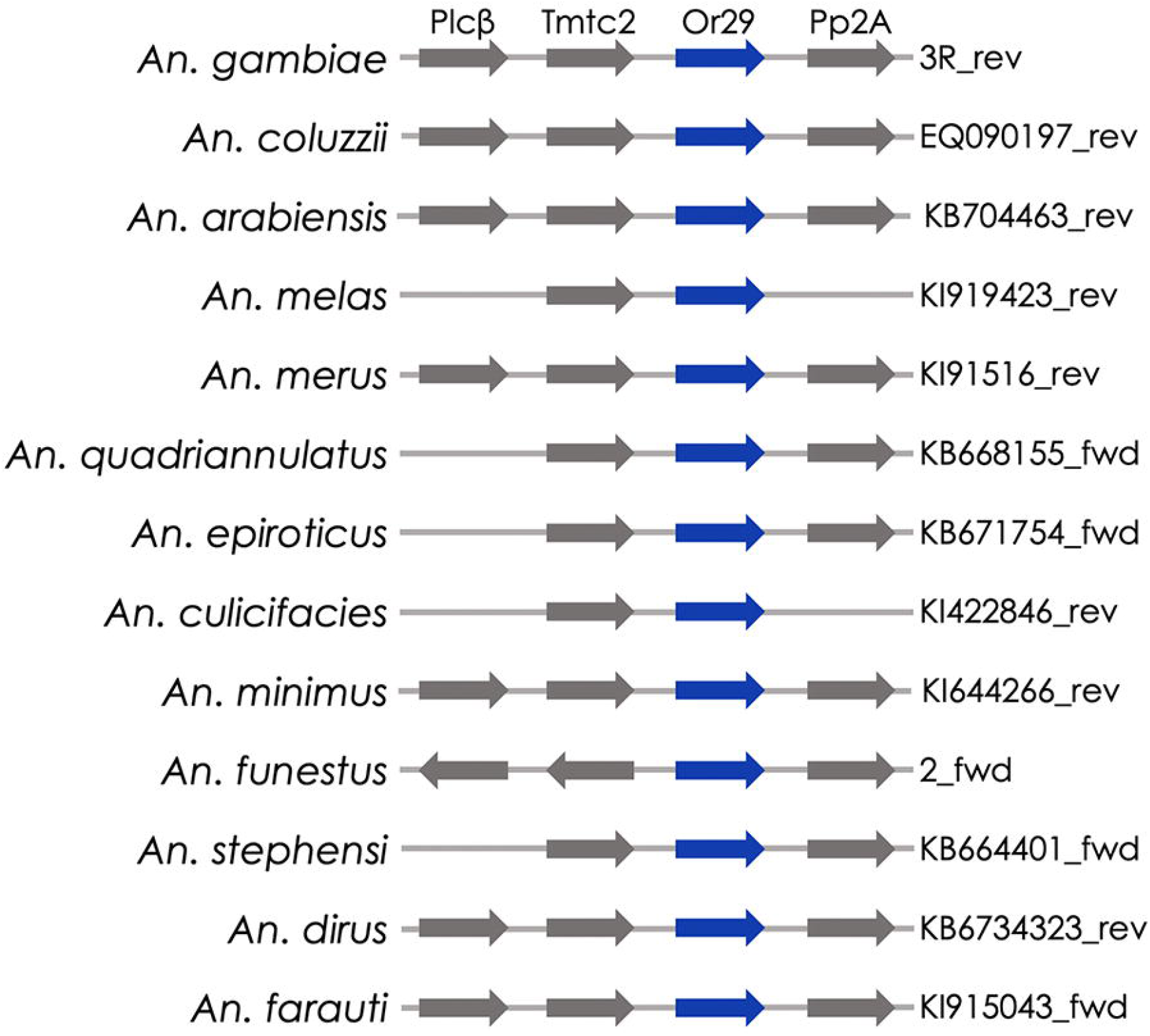
Microsyntenic relationships in anopheline genomes for Or29 orthologs. Relative positions of Or29 (blue) and conserved neighboring genes. Direction of transcription is indicated by arrows. Distances between genes are not drawn to scale. For *An. gambiae*, chromosome number and base pairing for the OR cluster are shown to the right. For all other species, chromosome or scaffold number is shown to the right of each genomic region, followed by the strand direction of either forward (fwd) or reverse (rev). A full listing of genes and abbreviations is provided in Supplemental Table S1.

**Fig. 3.**
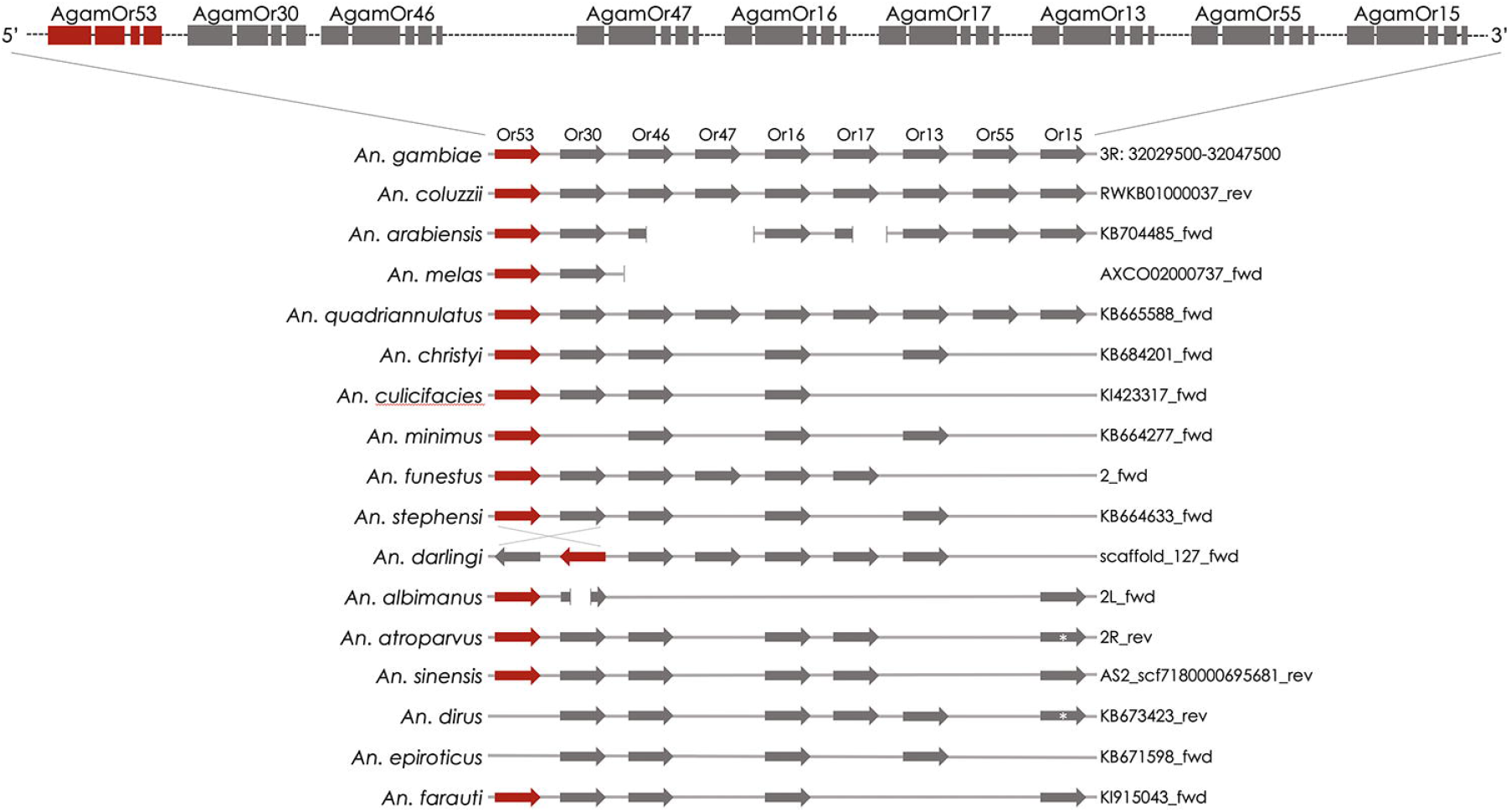
Microsyntenic relationships in anopheline genomes for Or53 orthologs and additional Ors. Exon/Intron structures of *An. gambiae* Or53 and linked Or cluster (Top). Relative positions of Or53 (red) and conserved neighboring genes. Direction of transcription is indicated by arrows. Distances between genes are not drawn to scale. For each species, chromosome or scaffold number is shown to the right of each genomic region, followed by the strand direction of either forward (fwd) or reverse (rev). A full listing of genes and abbreviations is provided in Supplemental Table S1.

**Fig. 4.**
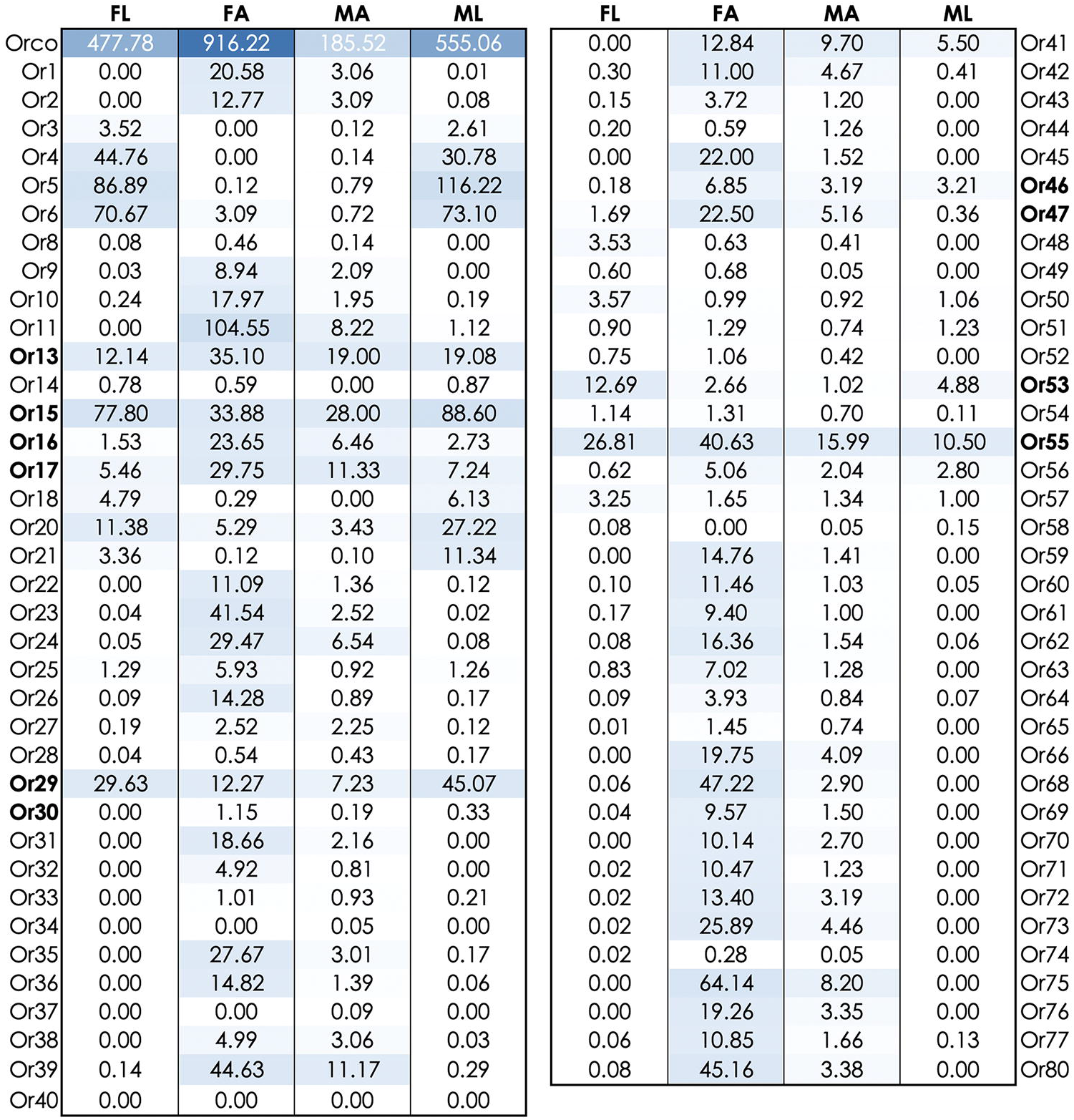
Odorant receptor (Or) expression in adult labellar lobes and antennae. Numbers represents average expression values given in Fragments Per Kilobase per Million (FPKM) for *An. coluzzii* labella (data from Saveer *et al*. 2018) or Reads Per Kilobase per Million (RPKM) for *An. gambiae* antennae (data from Pitts *et al*. 2011). Color intensity darkens with increasing expression values. Receptors indicated in bold are homologous genes shown in Figures 2 and 3. Abbreviations: FL, female labella; FA, female antennae; MA, male antennae; ML, male labella.

### Or29 and Or53 are tuned to linalool

Previously we characterized AgamOR29 as a linalool receptor, with apparent enantioselectivity (Huff and Pitts 2019). However, we did not investigate AgamOr29 sensitivity to the S-(+)-linalool isomer directly. In this study, we investigated the responsiveness of AgamOr29, AgamOr53, AsteOr29, and AsteOr53 to a panel of odorants, which included 119 organic compounds representing a range of chemical classes (Supplementary Table S2). We expressed the tuning receptors in combination with the corresponding coreceptors AgamOrco or AsteOrco, in *Xenopus laevis* oocytes and utilized two-electrode voltage clamping to record responses of the receptor complexes to blends of odorants (Supplementary Figure S1). Oocytes expressing functional receptor complexes were most strongly activated by Alcohol/Aldehyde blend 3, with at least 3-fold greater amplitude responses on average than any other odorant compound blend (Supplementary Figure S1). Oocytes expressing individual receptor complex subunits alone were unresponsive to any of the odorant blends (Supplementary Table S3). We next tested each of the individual chemicals that comprised Alcohol/Aldehyde blend 3 for each of the odorant receptor complexes. AgamOr29 and AsteOr29 in combination with their conspecific coreceptors showed the largest amplitude responses to (*S*)-(+)-linalool while AgamOr53 and AsteOr53 showed the largest amplitude responses to (*S*)-(+)-linalool with similar, albeit lower, responsiveness to the (*R*)-(–) enantiomer (Figure 5).

**Fig. 5.**
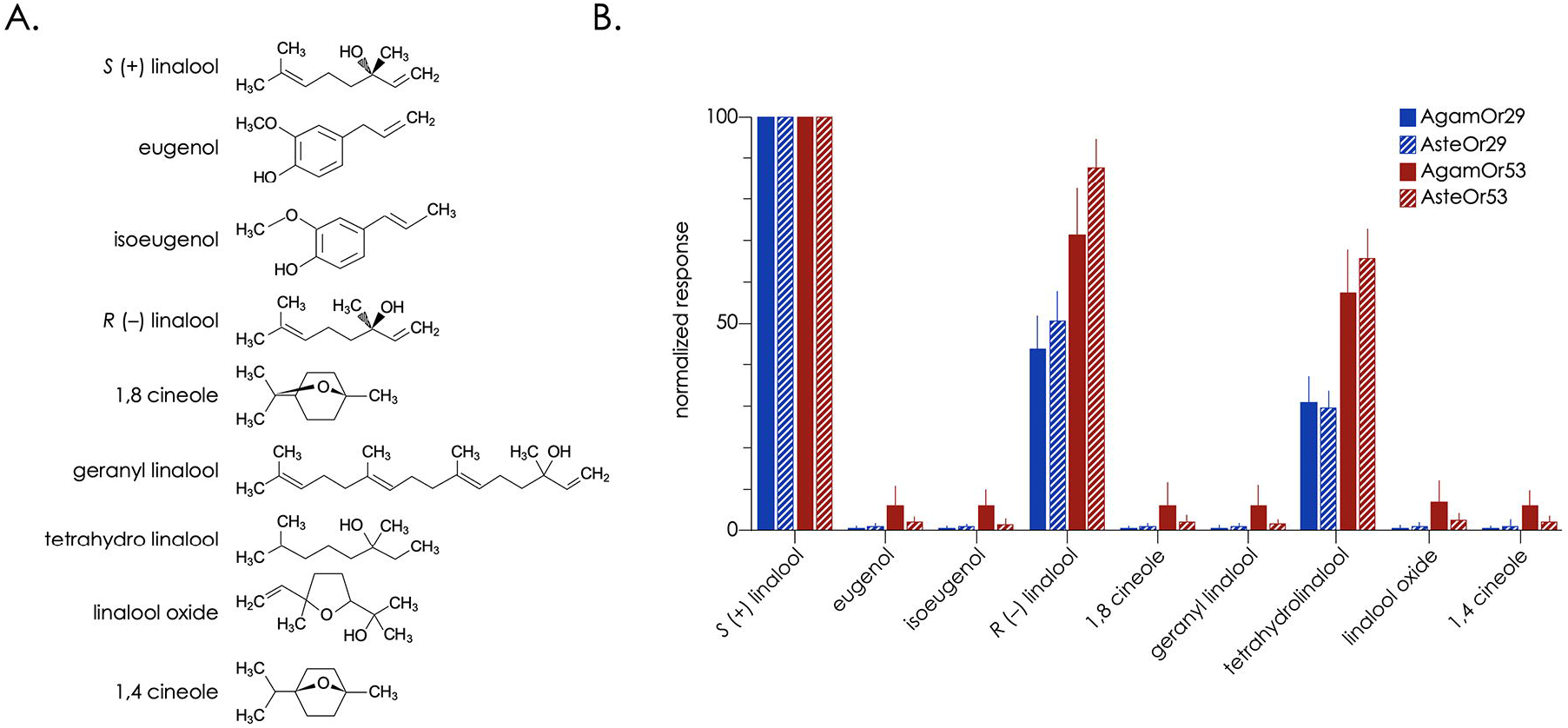
*An. gambiae* (filled) and *An. stephensi* (hashed) orthologs of Or29 (blue) and Or53 (red) are tuned to enantiomers of the monoterpene compound, linalool. (A) Chemical structures of the compounds used during the experiments are shown to the left. (B) Normalized mean current responses of oocytes (n = 7-10) coinjected with tuning receptor + conspecific Orco to individual alcohol compounds [10^−4^ M]. Standard errors are indicated in the positive direction only.

### Concentration dependency and receptor responses to linalool stereoisomers

A hallmark of receptor-ligand interactions is concentration dependency. In our functional assays, we challenged conspecific Orco/Or29 and Orco/Or53 receptor complexes to dilutions of both enantiomers of linalool across a broad range of concentrations, from 10^−7^ M to 10^−4^ M (Figure 6). The resulting electrophysiological responses were fitted to sigmoid curves and half-maximal effective concentration values (EC_50_) were calculated for isomers of linalool (Figure 6). The Or29 orthologs demonstrated two-fold lower EC_50_ values in response to the (*S*)-(+)-linalool isomer (Figure 6). Conversely, AsteOr53 produced a two-fold lower EC_50_ in response to the (*R*)-(–)-linalool, while the response of AgamOr53 was only moderately reduced (Figure 6).

**Fig. 6.**
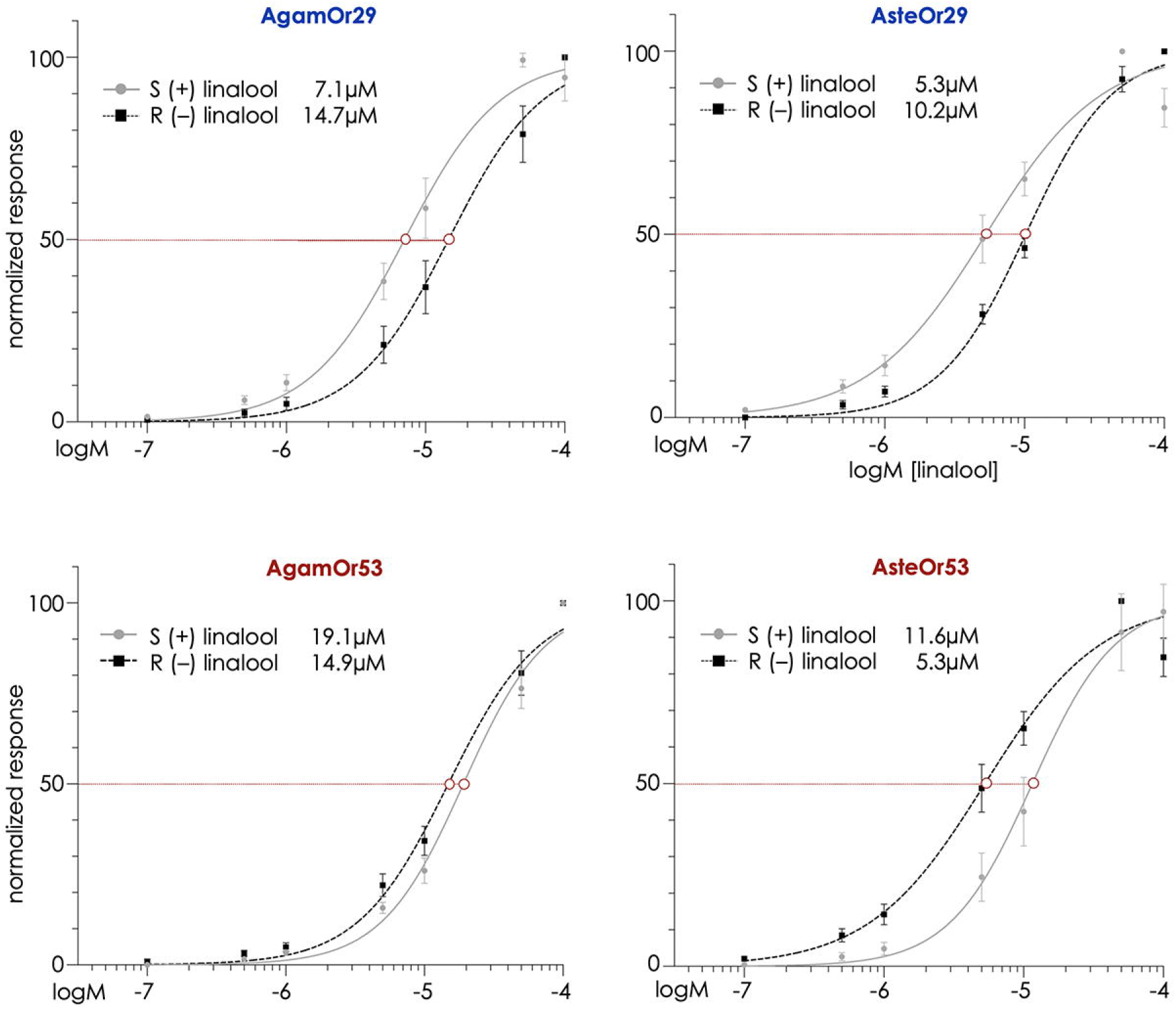
Concentration response curves of *An. gambiae* and *An. stephensi* orthologous odorant receptors to enantiomers of linalool. Normalized mean responses of oocytes injected with AgamOr29 + AgamOrco oocytes (top left) or AsteOr29 + AsteOrco (top right) to (S)-(+)-linalool (n = 10, gray circles) and (R)-(–)-linalool (n = 10, black squares). Normalized mean responses of oocytes injected with AgamOr53 + AgamOrco (lower left) or AsteOr53 + AsteOrco (lower right) to (S)-(+)-linalool (n = 10, gray circles) and (R)-(–)-linalool (n = 10, black squares). Estimated EC_50_ values in µM are provided for each CRC and the half maximal response is shown as a dashed red line.

## Discussion

Terpenes and terpenoids are emitted by many plant species and have profound effects on the chemical ecology of insects, acting as attractants and/or deterrents (Langenheim 1994; Degenhardt *et al*. 2003; Gillij *et al*. 2008; Muller *et al*. 2008). Many plants attract mosquitoes and extracts or synthetic blends derived from plant scent profiles are equally as attractive as the native odors (Gouagna *et al*. 2010; Nyasembe *et al*. 2012; Lutz *et al*. 2017). For example, *Culex pipiens* mosquitoes are attracted to floral extracts from *Asclepias syriaca* as well as a synthetic floral-odor blend mimicking the floral extract (Chaiphongpachara *et al*. 2018). Headspace volatiles of African plants, which include significant amounts of terpenoids, attract both sexes of *An. gambiae* (Nikbakhtzadeh *et al*. 2014), although linalool was not specifically identified in this study as a major constituent of the species examined. Nonetheless, linalool is a multifunctional monoterpene alcohol that may serve as a common pollinator attractant, being nearly ubiquitous in flowers (Crombie 1993; Knudsen and Tollsten 1993; Raguso 1999; Riffell *et al*. 2009). Linalool exists in two distinct forms in nature that are synthesized by enantiospecific biosynthetic enzymes (Ginglinger *et al*. 2013; Raguso 2016). Importantly, enantiomers of linalool can influence oviposition in insects, specifically in moths (van Schie *et al*. 2007; Reisenman *et al*. 2010b; Reisenman *et al*. 2013). One study on neuronal organization in *Manduca sexta* revealed a pair of unique glomeruli within female antennal lobes that exclusively receive inputs when antennae were stimulated with linalool (King *et al*. 2000). Follow up studies revealed projection neurons innervating female-specific glomeruli that demonstrated enantioselective responses to (R)-(−)- and (S)-(+)-linalool, which was not found in male antennal lobes (Reisenman *et al*. 2004).

Here we have shown that a pair of odorant receptors encoded within the genomes of both *An. gambiae* and *An. stephensi* are activated by enantiomers of linalool in the low micromolar concentration range (between 5 and 15_μ_M), consistent with the idea that it is the cognate odorant ligand for these receptors. All receptor combinations showed highest responses to the S-(+)-linalool isomer, indicating that this compound is the most efficacious (Figure 5). However, the potency of linalool enantiomers differed between the Or29 and Or53 orthologs (Figures 6). The (S)-(+) isomer produced a two-fold lower EC_50_ value for both AgamOr29 and AsteOr29, while the (R)-(−) isomer produced a two-fold lower EC_50_ value for AsteOr53. However, the apparent bias toward (R)-(−)-linalool was less pronounced for AgamOr53 (Figure 6).

The behavioral implications of having multiple receptors in Anophelines that are tuned to structurally similar stereoisomers is unclear. In the case of nectar feeding, one possibility is that racemic compounds such as linalool provides a more refined mechanism for discriminating preferred plant hosts. Linalool isomers are found in dramatically differing ratios across plant families and even within the same family (Ravid *et al*. 1997). Like other odors having stereoisomers, linalool enantiomers may be perceived by mosquitoes with opposing valencies; some isomers may elicit distinct behaviors, either attractive or repulsive, depending on blend composition and the physiological state of the animal (Omondi *et al*. 2019). Our tuning curves also reveal that tetrahydro linalool elicited a moderate, but repeatable response in both OR29 and Or53 orthologs, likely due to its structural similarity to linalool (Figure 5). This observation supports an odor coding mechanism whereby chemical ligands occupy an external binding pocket on their cognate receptor(s) and that compounds with similar structure can occupy the orthosteric site, albeit with lower binding affinities than the natural ligand (Saberi and Seyed-allaei 2016; Liang *et al*. 1998). Although not specifically investigated here, knowledge of stereoisomer selectivity, when combined with protein structural modeling and mutational analysis, can be used to investigate the potential interactions between ligands and amino acid residues of insect odorant receptors (Pellegrino *et al*. 2011; Leary *et al*. 2012; Hopf *et al*. 2015; Corcoran *et al*. 2018; de Mármol *et al*. 2021; Yuvaraj *et al*. 2021). The Anopheline Or29 and Or53 orthologs present an interesting opportunity for future in-depth studies of structure-activity relationships. We also acknowledge the possibility that additional odorant receptors in Anophelines may also be tuned to linalool, linalool derivatives, or other structurally related monoterpenes. In that case, unraveling the complex interplay between receptor activation in peripheral sensory neurons, plus the integration of signaling in the antennal lobe, will be crucial to understanding ultimate behavioral outputs.

In addition to its potential importance of in plant-seeking, linalool can also play a role in detection of blood meal hosts by vector mosquitoes. The skin microbiome can facilitate production and emanation of this compound and can provide a secondary behavioral contextual cue (Gallagher *et al*. 2008; Logan *et al*. 2008; Roodt *et al*. 2018; Omondi *et al*. 2019). The microbiome plays an important role in converting non-volatile skin compounds into VOCs (James *et al*. 2004a; James *et al*. 2004b). When skin bacteria are cultured on blood agar, they are attractive to *An. gambiae* in olfactometry experiments (Verhulst *et al*. 2009; Hill *et al*. 2009). The human skin microbiota and the volatiles that they emit can provide host seeking mosquitoes with a signal that a blood meal host is nearby (Meijerink *et al*. 2000; Verhulst *et al*. 2009). Perhaps bacterial derived skin volatiles and other semiochemicals including linalool can be modulated to increase the effectiveness of bait and kill pest management practices (Witzgall *et al*. 2010Kline 2007; Logan and Birkett 2007).

Further characterization of mosquito odorant receptors can inform these novel strategies by either elucidating the naturally evolved ligand-receptor combinations that may themselves offer the best targets for odor-based surveillance and trapping. Alternatively, additional agonists or antagonists of odorant receptors, especially those that hyperactivate or block the normal receptor function, could be utilized in “push-pull” vector control strategies (Tauxe et al 2013; Witzgall 2010). Pairs of closely related receptors that are biased toward opposing stereoisomers perhaps offer a unique opportunity to explore the behavioral outcomes in wild type or genetic knockout lines, particularly across species. Indeed, more comparative studies of insect chemoreceptors are needed to help inform our basic understanding of the molecular logic of odor coding. In this case, *An. gambiae* and *An. stephensi* are two of the most important malaria vectors,

## Acknowledgements

The authors thank Baylor University for research funding and core facility support. We thank Shan Ju Shih (Baylor University) for technical support. We also acknowledge Xenopus I Inc. (Dexter, MI) and Ecocyte Bioscience (Austin, TX) for supplying *Xenopus laevis* oocytes.

