## Supplementary figures and images for "Functional conservation of Anopheline linalool receptors through 100 million years of evolution"

### Supplementary Figure S1

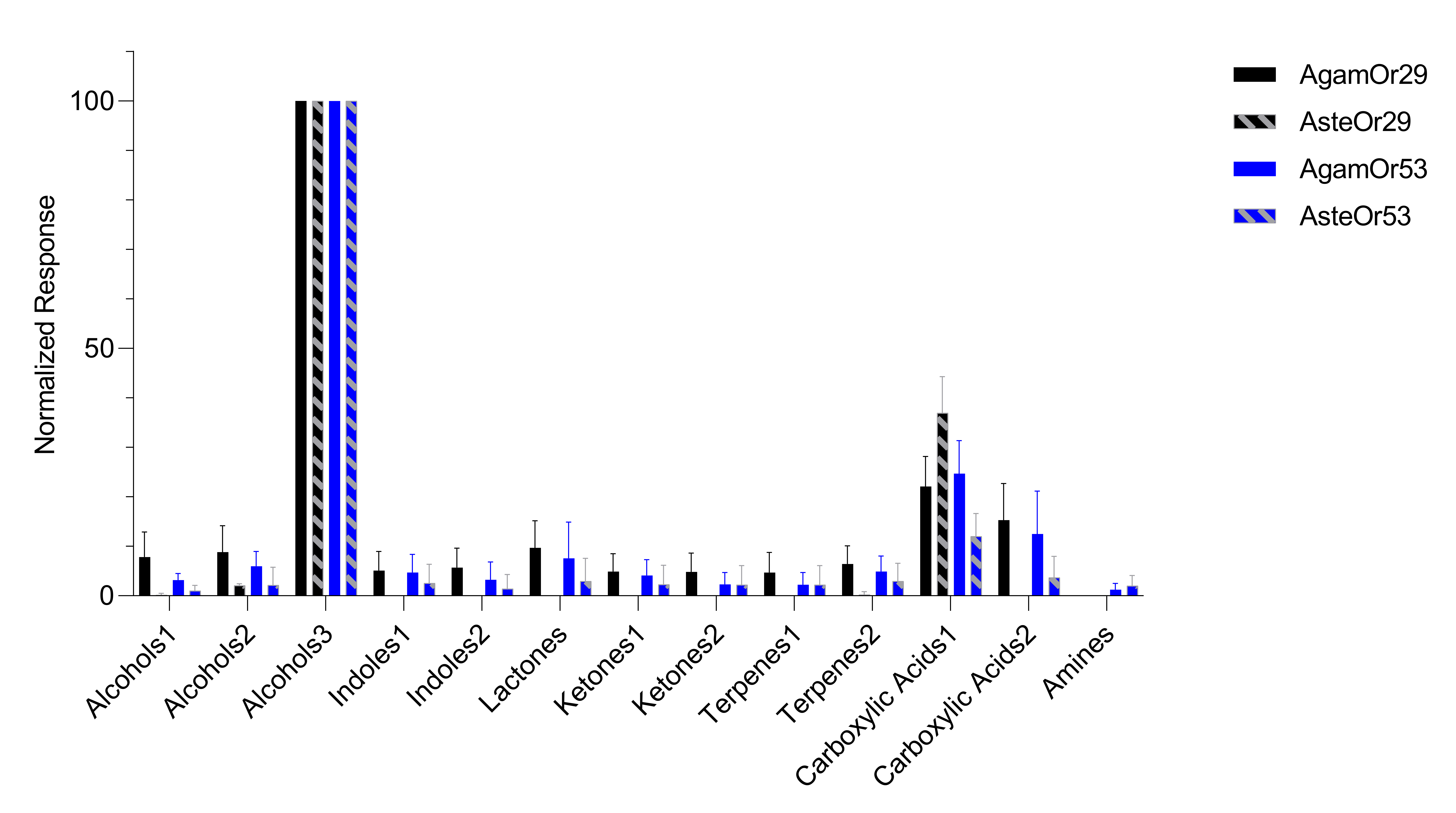
